# The Relationship between Body Mass Index and Poor Self-rated Health in the Korean Population

**DOI:** 10.1101/688994

**Authors:** Eun-Seok Sung, Chang Kyun Choi, Ji-An Jeong, Min-Ho Shin

**Affiliations:** Department of Public Health Graduate School, Chonnam National University, Gwangju, Republic of Korea; Department of Preventive Medicine, Chonnam National University Medical School, Hwasun, Republic of Korea

**Keywords:** Body Mass Index, Cross-Sectional Studies, Health Surveys

## Abstract

**Objective:** Several previous studies have evaluated associations between body mass index (BMI) and self-rated health (SRH); however, the results were inconsistent. This study aimed to examine the association between BMI and SRH in Korean adults.

**Methods:** The study was conducted in 214,997 adults who participated in the 2016 Korean Community Health Survey. Participants were categorized into four groups based on BMI: underweight (<18.5 kg/m^2^), normal weight (18.5–24.9 kg/m^2^), overweight (25.0–29.9 kg/m^2^), or obese (≥30.0 kg/m^2^). Multivariate Poisson regression analysis with sampling weights and robust variance estimators was performed to evaluate the relationship between BMI categories and poor SRH.

**Results:** There was a J-shaped association between BMI and poor SRH in both sexes, with the lowest risk observed in the normal weight group in both sexes. Compared with normal weight subjects, the age and lifestyle adjusted prevalence rate ratios for poor SRH were 1.61 (95% CI, 1.50–1.74) for underweight, 1.16 (95% CI, 1.11–1.21) for overweight, and 2.35 (95% CI, 2.13–2.58) for obese men; and 1.24 (95% CI, 1.17–1.32) for underweight, 1.26 (95% CI, 1.22–1.31) for overweight, and 1.77 (95% CI, 1.64–1.91) for obese women.

**Conclusions:** In a cross-sectional study using a nationally representative survey, there was a nonlinear relationship between BMI and poor SRH. This relationship was more prominent in men than in women. Prospective studies are needed to further clarify the relationship between BMI and SRH.

## Introduction

Obesity is an important risk factor in the chronic disease burden worldwide [1–3]. It is associated with increased morbidity and mortality of various chronic diseases [4]. Recently, underweight has also been associated with high all-cause and cardiovascular mortality [5–8]. However, it is unclear what health-related factors or diseases mediate this association [9]. Self-rated health (SRH) is an indicator of health condition that is a composite of physical, mental, and social well-being [10]. SRH is not only an index that reflects quality of life, but is also related to health conditions. The association of SRH with morbidity and mortality has been demonstrated not only in the general population but also in patients with various chronic diseases [11, 12]. In addition, SRH is an important prognostic predictor of diseases such as cancer and heart disease [13–15].

Several studies have explored the association between BMI and SRH [16–21]; however, the results have been inconsistent. Some studies have reported that higher BMI is associated with reduced SRH [19, 20]. Others have reported a U- or J-shaped association between BMI and poor SRH [16–18, 21, 22]. However, in previous studies, only the association between obesity and self-rated health was assessed, or the association between BMI and SRH was not properly assessed due to a lack of underweight subjects. Although there was one study in the Korean population, this one was based on a relatively small study population [23]. This study aimed to investigate the association between BMI and SRH in Korean adults, and to evaluate whether this association is modified by sex.

## Material and methods

### Subjects

This study used data from the 2016 Korean Community Health Survey (KCHS), which is a detailed survey of a representative sample of the Korean population aged 19 years or over conducted annually since 2008 to provide health statistics at the level of the municipality [24]. The target population for the KCHS is adults aged ≥ 19 years who live within jurisdiction of a community health center. The stratum was divided into two stages according to administrative unit (*Dong*, *Eup*, and *Myeon*) and housing unit (apartments and houses), and the smallest administrative district unit (*Tong*, *Ban*, and *Lee*) was selected as the primary sampling unit of the stratum through probability proportionate sampling. Information was gathered through face-to-face interviews conducted by a trained interviewer. The 2016 KCHS included 228,452 subjects, but 214,997 subjects (99,930 men and 115,067 women) were included in this study due to missing values. This study was approved by the Institutional Review Board of the Korea Centers for Disease Control and Prevention (KCDC).

### Body Mass Index

BMI was based on self-reported height and weight and calculated as weight in kilograms divided by height in meters squared. Participants were categorized into four groups based on BMI: underweight (<18.5 kg/m^2^), normal weight (18.5–24.9 kg/m^2^), overweight (25.0–29.9 kg/m^2^), or obese (≥30.0 kg/m^2^). Normal weight was chosen as the reference category.

### Self-Rated Health

SRH was assessed with the question, “How do you rate your general health status?”, rated on a 5-point scale. SRH was dichotomized as poor (reported as “poor” or “very poor”) or good (reported as “very good,” “good,” or “moderate”).

### Covariates

Lifestyle, socioeconomic status, and comorbidities were investigated by interview. Smoking history was coded as non-smoker, former smoker, or current smoker. Alcohol intake was coded as drinker or non-drinker. Socioeconomic variables considered in this study included marital status (single, married, or divorced/bereaved/separated), educational attainment (uneducated, elementary school, middle school, high school, or college or higher), rural residency, and monthly household income (≤1.00 million, 1.01–2.00 million, 2.01–3.00 million, 3.01–4.00 million, or ≥4.01 million South Korean won). Physical activity was coded as whether subjects engaged in moderate or vigorous physical activity. Moderate physical activity was defined as moderate intense physical activity (e.g., swimming at a slow pace, table tennis, badminton, tennis doubles) for more than 5 days per week for 30 min or more. Vigorous physical activity was defined as vigorous intense physical activity (e.g., swimming at a fast pace, climbing, cycling, squash, tennis singles) for more than 3 days per week for 20 min or more. Co-morbidities (if respondents had doctor-diagnosed hypertension, diabetes, stroke, coronary heart disease, or arthritis) were also considered.

### Statistical analysis

The data were analyzed separately for each sex. Tests for trends across BMI categories were performed using linear regression. The relationship between BMI and SRH was assessed using multivariate Poisson regression analysis with sampling weights and robust variance estimators. The prevalence rate ratio (PRR) and corresponding 95% confidence intervals (CIs) were reported. Restricted cubic splines were used to model the nonlinear relationship between BMI and SRH using knots at 5th, 50th, and 95th percentile of BMI. Stata version 14.0 (Stata Corp, College Station, TX, USA) was used for the statistical analyses. Statistical significance was defined as a *P*-value < 0.05.

## Results

The study population included 99,930 men and 115,067 women. Baseline characteristics according to BMI categories are presented in Tables 1 and 2. The prevalence of underweight was 2.7% in men and 7.0% in women, while the prevalence of obesity was 3.6% in men and 2.5% in women. In both sexes, subjects with higher BMI were poor SRH, older, married, had higher income and education levels, lived in cities, engaged in more physical activity, and had higher alcohol consumption. In addition, they had higher rates of hypertension and diabetes, but lower rates of stroke, coronary heart disease, and arthritis. Among female participants, subjects with higher BMI were older, less likely to be married, had lower income and education levels, lived in rural areas, engaged in more physical activity, and had higher alcohol consumption. In addition, they had higher rates of hypertension, diabetes, arthritis, stroke, and coronary artery disease. Current smokers were more common among underweight and obese subjects in both women and men. Among the women, the underweight and obese groups had lower physical activity. To control for confounding effects, sensitivity analysis was performed only for non-smokers or subjects without arthritis, coronary heart disease, or stroke. However, the results of the sensitivity analysis were not significantly different from those of the existing analysis, and are not shown in the results.

**Table 1.**
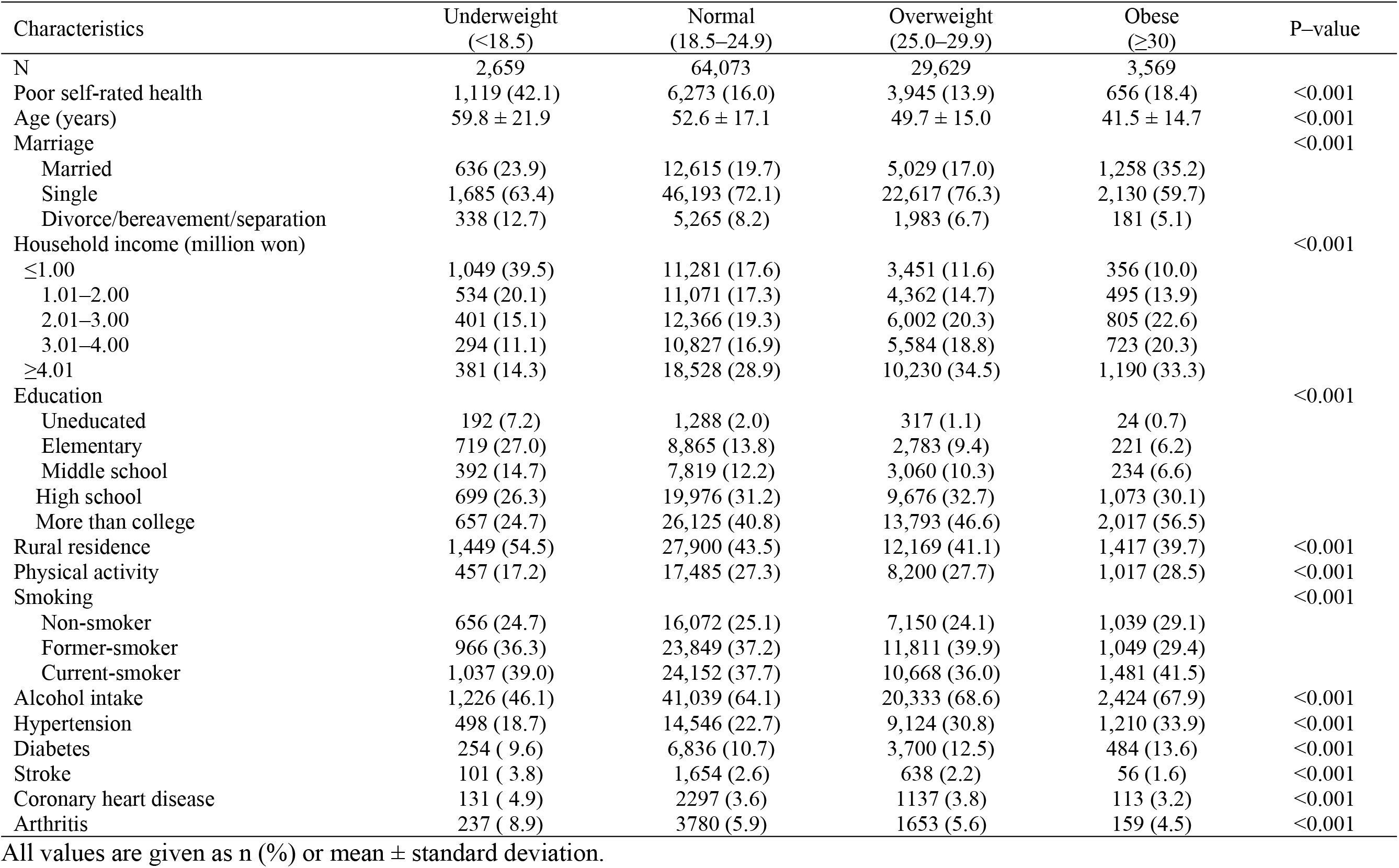
Baseline characteristics according to body mass index (kg/m^2^) categories in men.

**Table 1.** Baseline characteristics according to body mass index (kg/m^2^) categories in women.

| Characteristics | Underweight<br>($<18.5$) | Normal<br>($18.5\text{--}24.9$) | Overweight<br>($25.0\text{--}29.9$) | Obese<br>($\geq 30$) | P-value |
| --- | --- | --- | --- | --- | --- |
| N | 7,997 | 81,929 | 22,225 | 2,916 |  |
| Poor self-rated health | 2,081 (26.0) | 16,365 (20.0) | 6,344 (28.5) | 1,063 (36.5) | $<0.001$ |
| Age (years) | $47.6 \pm 21.8$ | $51.8 \pm 17.1$ | $56.2 \pm 14.8$ | $52.3 \pm 16.5$ | $<0.001$ |
| Marriage | | | | | $<0.001$ |
| Married | 2,403 (30.0) | 10,988 (13.4) | 1,430 (6.4) | 392 (13.4) |  |
| Single | 3,818 (47.7) | 54,121 (66.1) | 15,403 (69.3) | 1,842 (63.2) |  |
| Divorce/bereavement/separation | 1,776 (22.2) | 16,820 (20.5) | 5,392 (24.3) | 682 (23.4) |  |
| Household income (million won) | | | | | $<0.001$ |
| $\leq 1.00$ | 1,863 (23.3) | 16,065 (19.6) | 5,311 (23.9) | 715 (24.5) | |
| 1.01–2.00 | 1,034 (12.9) | 13,111 (16.0) | 4,314 (19.4) | 561 (19.2) |  |
| 2.01–3.00 | 1,257 (15.7) | 14,324 (17.5) | 4,153 (18.7) | 579 (19.9) |  |
| 3.01–4.00 | 1,194 (14.9) | 13,332 (16.3) | 3,355 (15.1) | 465 (15.9) |  |
| $\geq 4.01$ | 2,649 (33.1) | 25,097 (30.6) | 5,092 (22.9) | 596 (20.4) | |
| Education | | | | | $<0.001$ |
| Uneducated | 1,025 (12.8) | 6,272 (7.7) | 1,718 (7.7) | 223 (7.6) |  |
| Elementary | 1,194 (14.9) | 15,045 (18.4) | 6,190 (27.9) | 730 (25.0) |  |
| Middle school | 367 (4.6) | 8,667 (10.6) | 3,596 (16.2) | 414 (14.2) |  |
| High school | 1,497 (18.7) | 22,857 (27.9) | 6,262 (28.2) | 904 (31.0) |  |
| More than college | 3,914 (48.9) | 29,088 (35.5) | 4,459 (20.1) | 645 (22.1) |  |
| Rural residence | 3,131 (39.2) | 33,079 (40.4) | 10,246 (46.1) | 1,280 (43.9) | $<0.001$ |
| Physical activity | 1,159 (14.5) | 16,017 (19.5) | 4,309 (19.4) | 490 (16.8) | $<0.001$ |
| Smoking | | | | | $<0.001$ |
| Non-smoker | 7,331 (91.7) | 77,685 (94.8) | 20,962 (94.3) | 2,638 (90.5) |  |
| Former-smoker | 258 (3.2) | 1,931 (2.4) | 632 (2.8) | 135 (4.6) |  |
| Current-smoker | 408 (5.1) | 2,313 (2.8) | 631 (2.8) | 143 (4.9) |  |
| Drinking | 2,534 (31.7) | 25,302 (30.9) | 5,670 (25.5) | 788 (27.0) | $<0.001$ |
| Hypertension | 1,169 (14.6) | 17,406 (21.2) | 8,611 (38.7) | 1,338 (45.9) | $<0.001$ |
| Diabetes | 394 (4.9) | 6,387 (7.8) | 3,211 (14.4) | 594 (20.4) | $<0.001$ |
| Stroke | 125 (1.6) | 1,251 (1.5) | 543 (2.4) | 81 (2.8) | $<0.001$ |
| Coronary heart disease | 179 (2.2) | 1,942 (2.4) | 898 (4.0) | 142 (4.9) | $<0.001$ |
| Arthritis | 1081 (13.5) | 13,178 (16.1) | 5,909 (26.6) | 830 (28.5) | $<0.001$ |

All values are given as n (%) or mean ± standard deviation.

Table 3 presents the PRRs for poor SRH according to BMI categories. There was a nonlinear relationship between BMI and poor SRH. The age-adjusted PRR for poor SRH across BMI levels (<18.5, 25–29.9, and ≥30 kg/m^2^) was 1.99 (95% CI, 1.85–2.15), 1.09 (95% CI, 1.04–1.14), and 2.48 (95% CI, 2.26–2.73), respectively, for men; and 1.33 (95% CI, 1.26–1.41), 1.34 (95% CI, 1.29–1.39), and 2.04 (95% CI, 1.89–2.20), respectively, for women. After further adjustment for demographic, socioeconomic, and lifestyle characteristics, this U-shaped association was slightly attenuated but remained statistically significant in both sexes. In model 3, the adjusted PRR for BMI <18.5 kg/m^2^ was 1.61 (95% CI, 1.50–1.74), that for BMI 25.0–29.9 kg/m^2^ was 1.16 (95% CI, 1.11–1.21), and that for BMI ≥30 kg/m^2^ was 2.35 (95% CI, 2.13–2.58) for men; and the corresponding values were 1.24 (95% CI, 1.17–1.32), 1.26 (95% CI, 1.22–1.31), and 1.77 (95% CI, 1.64–1.91), respectively, for women. With further adjustment for comorbidities, the PRRs of overweight and obesity were attenuated, whereas the PRR for underweight was strengthened. In model 4, the PRR for underweight was 1.83 (95% CI, 1.69–1.98), that for overweight was 1.04 (95% CI, 1.00–1.09), and that for obese was 1.90 (95% CI, 1.73–2.09) in men, and the corresponding values were 1.32 (95% CI, 1.25–1.40), 1.14 (95% CI, 1.10–1.18), and 1.41 (95% CI, 1.30–1.52), respectively, for women. The interaction by sex was significant (*P* for interaction < 0.001).

**Table 3.**
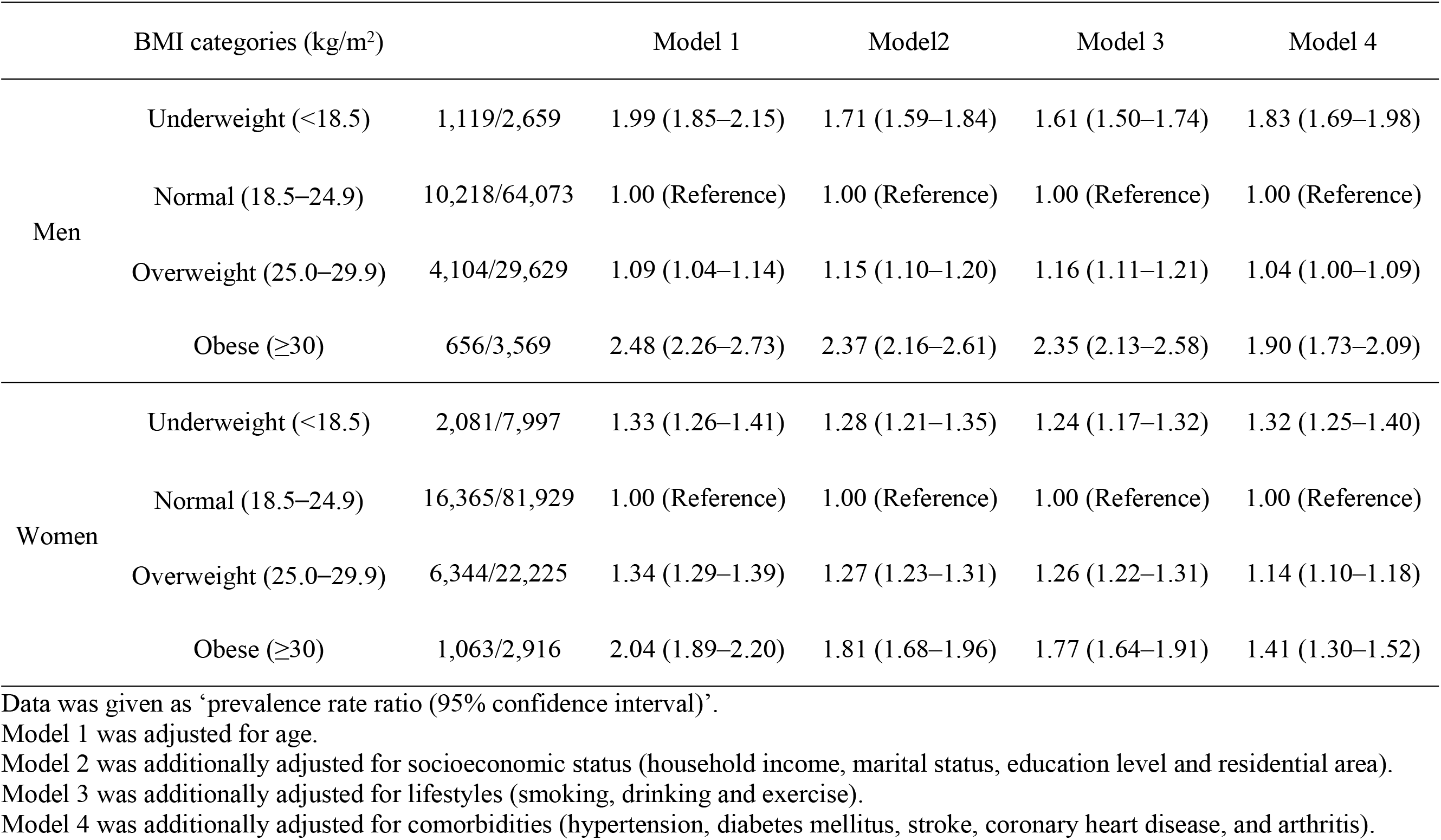
Association between body mass index and poor self-rated health.

Figure 1 and 2 presented the adjusted prevalence of poor SRH from restricted cubic spline models in both sexes. In both sexes, there was a J-shaped association between BMI and poor SRH, with the lowest risk observed with a BMI of 23–24.9 kg/m^2^ in both men and women. This association was more prominent in men than in women.

**Fig 1.**
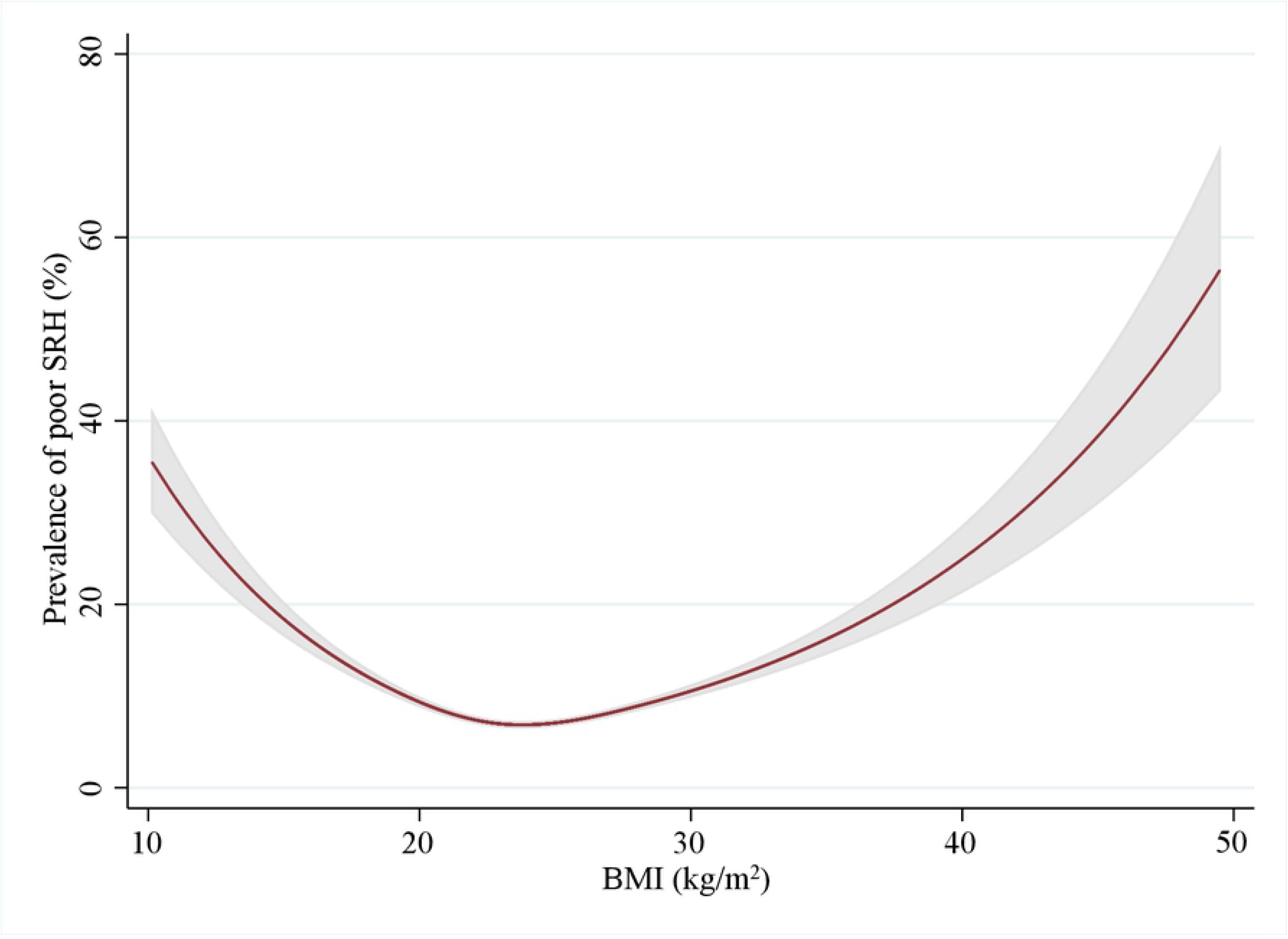
Adjusted restricted cubic splines for the association between body mass index and poor self-rated health in men. Age, lifestyle, socioeconomic status, and comorbidities were adjusted. Shaded areas indicated 95% confidence intervals.

**Fig 2.**
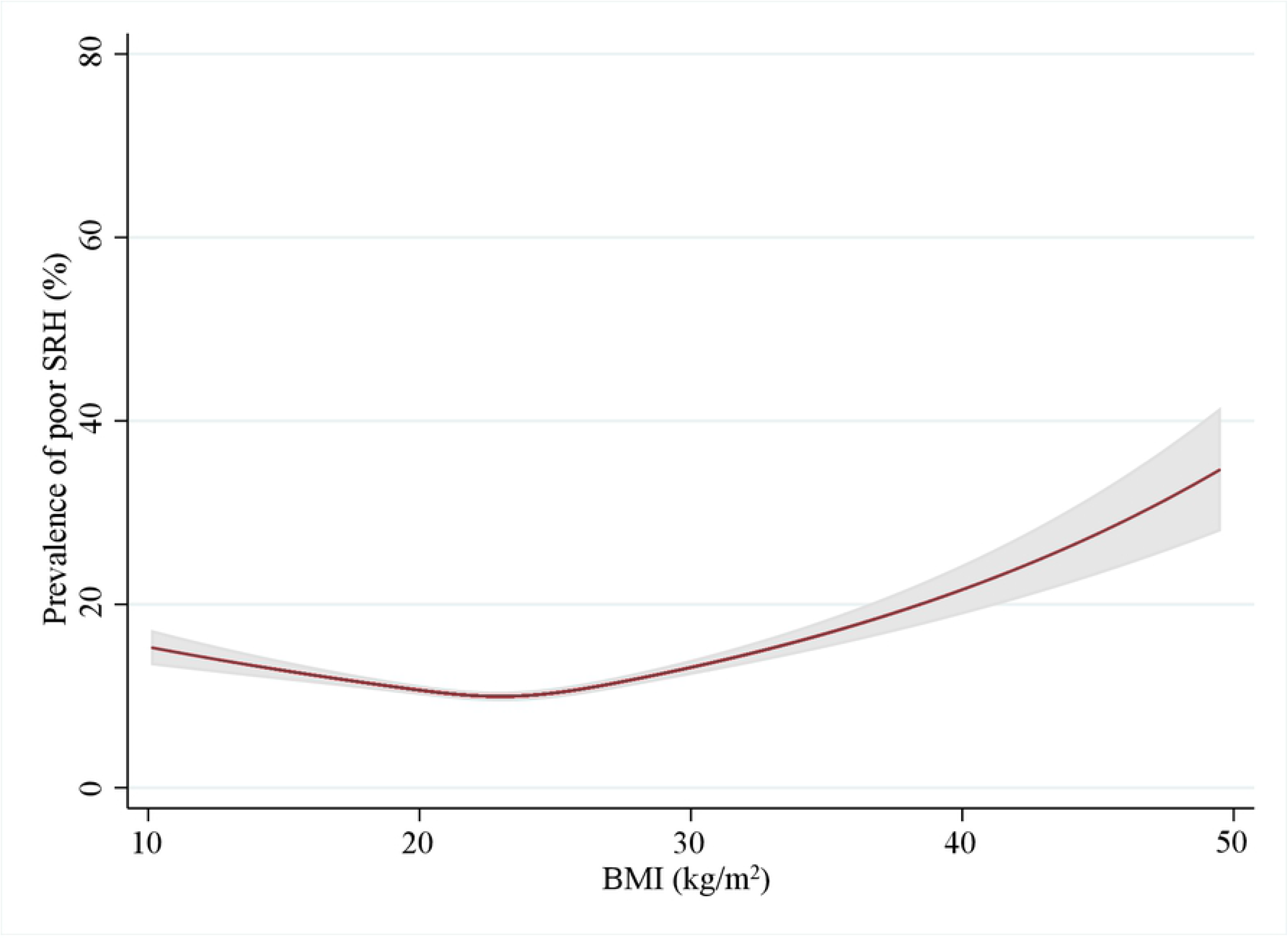
Adjusted restricted cubic splines for the association between body mass index and poor self-rated health in women. Age, lifestyles, socioeconomic status, and comorbidities were adjusted. Shaded areas indicated 95% confidence intervals.

## Discussion

In this cross-sectional study using a nationwide sample, both obesity and underweight were associated with an increased propensity for poor SRH, even after adjusting for demographic, socioeconomic, and lifestyle variables. Furthermore, a J-shaped association between BMI and poor SRH was more prominent in men than in women. This study is the first to assess the association between BMI and SRH in a national sample from a community health survey.

There was a J-shaped relationship between BMI and SRH in Korean adults, and this relationship was more prominent in men than in women. Except for studies in which the relationship with underweight was not evaluated [19, 20], previous studies have reported U- or J-shaped relationships but have been inconsistent regarding sex-specific relationships [17, 18, 21, 22]. Heo et al. reported a J-shaped relationship between BMI and health-related quality of life [21]. Similar to the present study, when adjusted for comorbidities, the association between underweight and poor SRH was strengthened, while the associations between overweight and obesity with poor SRH were attenuated. In the Canadian Community Health Survey [17], BMI showed a U-shaped association with poor SRH in both sexes. In a study based on the National Health Interview Survey from 1997 to 2005 [18] and a study based on the World Health Survey (2002–2004) [22], the association between underweight with poor SRH was more evident in men, whereas the association between obesity with poor SRH was more evident in women. However, the Sault Antenatal Care Programme in Sweden [16], found a U-shaped relationship in women, but a linear relationship in men. This study was conducted in expectant parents, rather than in the general population, and was a small study with fewer than 1,000 subjects, which may have resulted in differences from the results of previous studies. There was one study based on Korean. Although direct comparison is difficult because of differences in study design, the result of previous study were along the same line as our results. Noh et al. assessed the relationship between BMI categorized as WHO Asian classification and SRH scores, and SRH scores was the highest in the normal group (18.5–22.9 kg/m^2^) [23]. Compared to normal group, the odd ratio (OR) of poor SRH score was 1.60 in underweight, 1.08 in overweight, 1.72 in obese, and 3.97 in severe obese.

The sex difference in the association between BMI and SRH may be partly due to the sex difference in body fat distribution. At the same BMI, women tend to have higher body fat than men across all ages [25, 26]. On the other hand, visceral adipose tissue increases more in men as BMI increases in those under age 40, whereas the association between visceral adipose tissue and BMI does not differ according to sex in those aged 40 or older [26]. Unfortunately, because this study did not measure total body fat or lean body mass, whether the relationship between BMI and SRH varies depending on differences in fat distribution cannot be assessed.

The underlying mechanism of the relationship between underweight and poor SRH can be explained as follows. First, underweight is associated with sarcopenia [27]. Subjects with sarcopenia or decreased muscle mass have lower exercise capacity and lower levels of physical activity, both of which are associated with poor SRH. Second, the relationship between underweight and poor SRH may be a reverse causation. Hulman et al. reported that weight loss was greater in subjects with poor SRH than with good SRH [28].

The relationship between obesity and poor SRH may be mediated by obesity-related comorbidities. Numerous studies have reported causal relationships between obesity and cardiovascular diseases and cancer [29–31], which can lead to poor SRH and poor quality of life [32]. In our study, after adjusting for comorbidities, the strength of the relationship decreased by more than 30%. Additionally, obesity is associated with reduced physical activity and lower exercise capacity, and both conditions are associated with poor SRH.

The strength of this study is the large, nationally representative sample size. However, the study has some limitations. First, the design of our study is cross-sectional, which leads to difficulty in assessing causal relationships. Second, BMI was calculated using self-reported height and weight. However, there was a high correlation coefficient between measured BMI and self-reported BMI in the 2016 KCHS (r = 0.92) [33]. Third, other indices reflecting adiposity, such as waist circumference, total body fat, and subcutaneous abdominal adipose tissue mass, were not included in this study, and further evaluation using these indices is needed.

## Conclusion

In conclusion, there was a J-shaped relationship between BMI and poor SRH in a nationwide sample of Korean adults. These results suggest that both underweight and overweight are risk factors for poor SRH. However, prospective studies are needed to clarify the causal relationship between these two variables.

## Acknowledgements

This manuscript is a revision of the first author’s PhD thesis from Chonnam National University.

## References

1. Quesenberry CP, Jr., Caan B, Jacobson A. Obesity, health services use, and health care costs among members of a health maintenance organization. Archives of internal medicine. 1998;158(5):466–72. PubMed PMID: 9508224.

2. Visscher TL, Seidell JC. The public health impact of obesity. Annual review of public health. 2001;22:355–75. doi: 10.1146/annurev.publhealth.22.1.355. PubMed PMID: 11274526.

3. Flynn MA, McNeil DA, Maloff B, Mutasingwa D, Wu M, Ford C, et al. Reducing obesity and related chronic disease risk in children and youth: a synthesis of evidence with ‘best practice’ recommendations. Obesity reviews : an official journal of the International Association for the Study of Obesity. 2006;7 Suppl 1:7–66. doi: 10.1111/j.1467-789X.2006.00242.x. PubMed PMID: 16371076.

4. Abdelaal M, le Roux CW, Docherty NG. Morbidity and mortality associated with obesity. Ann Transl Med. 2017;5(7):161. Epub 2017/05/10. doi: 10.21037/atm.2017.03.107. PubMed PMID: 28480197; PubMed Central PMCID: PMCPMC5401682.

5. Flegal KM, Graubard BI, Williamson DF, Gail MH. Cause-specific excess deaths associated with underweight, overweight, and obesity. JAMA. 2007;298(17):2028–37. Epub 2007/11/08. doi: 10.1001/jama.298.17.2028. PubMed PMID: 17986696.

6. Ozaltin E, Hill K, Subramanian SV. Association of maternal stature with offspring mortality, underweight, and stunting in low- to middle-income countries. Jama. 2010;303(15):1507–16. doi: 10.1001/jama.2010.450. PubMed PMID: 20407060; PubMed Central PMCID: PMC3100588.

7. Lange BJ, Gerbing RB, Feusner J, Skolnik J, Sacks N, Smith FO, et al. Mortality in overweight and underweight children with acute myeloid leukemia. Jama. 2005;293(2):203–11. doi: 10.1001/jama.293.2.203. PubMed PMID: 15644547.

8. Visscher TL, Seidell JC, Menotti A, Blackburn H, Nissinen A, Feskens EJ, et al. Underweight and overweight in relation to mortality among men aged 40-59 and 50-69 years: the Seven Countries Study. American journal of epidemiology. 2000;151(7):660–6. PubMed PMID: 10752793.

9. Roh L, Braun J, Chiolero A, Bopp M, Rohrmann S, Faeh D, et al. Mortality risk associated with underweight: a census-linked cohort of 31,578 individuals with up to 32 years of follow-up. BMC Public Health. 2014;14:371. Epub 2014/04/18. doi: 10.1186/1471-2458-14-371. PubMed PMID: 24739374; PubMed Central PMCID: PMCPMC4021191.

10. Mossey JM, Shapiro E. Self-rated health: a predictor of mortality among the elderly. American journal of public health. 1982;72(8):800–8. PubMed PMID: 7091475; PubMed Central PMCID: PMC1650365.

11. Idler EL, Benyamini Y. Self-rated health and mortality: a review of twenty-seven community studies. Journal of health and social behavior. 1997;38(1):21–37. PubMed PMID: 9097506.

12. Heidrich J, Liese AD, Lowel H, Keil U. Self-rated health and its relation to all-cause and cardiovascular mortality in southern Germany. Results from the MONICA Augsburg cohort study 1984-1995. Annals of epidemiology. 2002;12(5):338–45. PubMed PMID: 12062922.

13. Osoba D. What has been learned from measuring health-related quality of life in clinical oncology. European journal of cancer. 1999;35(11):1565–70. PubMed PMID: 10673963.

14. Gill TM, Feinstein AR. A critical appraisal of the quality of quality-of-life measurements. Jama. 1994;272(8):619–26. PubMed PMID: 7726894.

15. Coates A, Porzsolt F, Osoba D. Quality of life in oncology practice: prognostic value of EORTC QLQ-C30 scores in patients with advanced malignancy. European journal of cancer. 1997;33(7):1025–30. PubMed PMID: 9376182.

16. Eurenius E, Lindkvist M, Sundqvist M, Ivarsson A, Mogren I. Maternal and paternal self-rated health and BMI in relation to lifestyle in early pregnancy: the Salut Programme in Sweden. Scand J Public Health. 2011;39(7):730–41. Epub 2011/09/21. doi: 10.1177/1403494811418279. PubMed PMID: 21930619.

17. Herman KM, Hopman WM, Rosenberg MW. Self-rated health and life satisfaction among Canadian adults: associations of perceived weight status versus BMI. Quality of life research : an international journal of quality of life aspects of treatment, care and rehabilitation. 2013;22(10):2693–705. doi: 10.1007/s11136-013-0394-9. PubMed PMID: 23539466.

18. Imai K, Gregg EW, Chen YJ, Zhang P, de Rekeneire N, Williamson DF. The association of BMI with functional status and self-rated health in US adults. Obesity (Silver Spring). 2008;16(2):402–8. doi: 10.1038/oby.2007.70. PubMed PMID: 18239651.

19. Okosun IS, Choi S, Matamoros T, Dever GE. Obesity is associated with reduced self-rated general health status: evidence from a representative sample of white, black, and Hispanic Americans. Prev Med. 2001;32(5):429–36. Epub 2001/05/02. doi: 10.1006/pmed.2001.0840. PubMed PMID: 11330993.

20. Prosper MH, Moczulski VL, Qureshi A. Obesity as a predictor of self-rated health. Am J Health Behav. 2009;33(3):319–29. Epub 2008/12/10. PubMed PMID: 19063653.

21. Heo M, Allison DB, Faith MS, Zhu S, Fontaine KR. Obesity and quality of life: mediating effects of pain and comorbidities. Obesity research. 2003;11(2):209–16. doi: 10.1038/oby.2003.33. PubMed PMID: 12582216.

22. Wang A, Arah OA. Body Mass Index and Poor Self-Rated Health in 49 Low-Income and Middle-Income Countries, By Sex, 2002-2004. Prev Chronic Dis. 2015;12:E133. Epub 2015/08/21. doi: 10.5888/pcd12.150070. PubMed PMID: 26292064; PubMed Central PMCID: PMCPMC4556100.

23. Fürnsinn C, Noh J-W, Kim J, Yang Y, Park J, Cheon J, et al. Body mass index and self-rated health in East Asian countries: Comparison among South Korea, China, Japan, and Taiwan. Plos One. 2017;12(8). doi: 10.1371/journal.pone.0183881.

24. Kang YW, Ko YS, Kim YJ, Sung KM, Kim HJ, Choi HY, et al. Korea Community Health Survey Data Profiles. Osong Public Health Res Perspect. 2015;6(3):211–7. Epub 2015/10/03. doi: 10.1016/j.phrp.2015.05.003. PubMed PMID: 26430619; PubMed Central PMCID: PMCPMC4551141.

25. Gallagher D, Visser M, Sepulveda D, Pierson RN, Harris T, Heymsfield SB. How useful is body mass index for comparison of body fatness across age, sex, and ethnic groups? Am J Epidemiol. 1996;143(3):228–39. Epub 1996/02/01. PubMed PMID: 8561156.

26. Camhi SM, Bray GA, Bouchard C, Greenway FL, Johnson WD, Newton RL, et al. The relationship of waist circumference and BMI to visceral, subcutaneous, and total body fat: sex and race differences. Obesity (Silver Spring). 2011;19(2):402–8. Epub 2010/10/16. doi: 10.1038/oby.2010.248. PubMed PMID: 20948514; PubMed Central PMCID: PMCPMC3960785.

27. Lauretani F, Russo CR, Bandinelli S, Bartali B, Cavazzini C, Di Iorio A, et al. Age-associated changes in skeletal muscles and their effect on mobility: an operational diagnosis of sarcopenia. Journal of Applied Physiology. 2003;95(5):1851–60. doi: 10.1152/japplphysiol.00246.2003.

28. Vinciguerra M, Hulman A, Ibsen DB, Laursen ASD, Dahm CC. Body mass index trajectories preceding first report of poor self-rated health: A longitudinal case-control analysis of the English Longitudinal Study of Ageing. Plos One. 2019;14(2). doi: 10.1371/journal.pone.0212862.

29. O’Rourke RW. Obesity and Cancer. Metabolic Syndrome and Diabetes 2016. p. 111–23.

30. Piche ME, Poirier P, Lemieux I, Despres JP. Overview of Epidemiology and Contribution of Obesity and Body Fat Distribution to Cardiovascular Disease: An Update. Prog Cardiovasc Dis. 2018;61(2):103–13. Epub 2018/07/03. doi: 10.1016/j.pcad.2018.06.004. PubMed PMID: 29964067.

31. Calle EE, Rodriguez C, Walker-Thurmond K, Thun MJ. Overweight, obesity, and mortality from cancer in a prospectively studied cohort of U.S. adults. N Engl J Med. 2003;348(17):1625–38. Epub 2003/04/25. doi: 10.1056/NEJMoa021423. PubMed PMID: 12711737.

32. Ko H-Y, Lee J-K, Shin J-Y, Jo E. Health-Related Quality of Life and Cardiovascular Disease Risk in Korean Adults. Korean Journal of Family Medicine. 2015;36(6). doi: 10.4082/kjfm.2015.36.6.349.

33. Jeong J-y, Kim D-H, Kim K-Y, Ryu SY, Lee S-Y, Park YS. Accuracy of Self-reported Height, Weight and Body Mass Index in Community Health Survey in South Korea. Journal of Health Informatics and Statistics. 2017;42(3):241–9. doi: 10.21032/jhis.2017.42.3.241.

